# A novel thermostable and processive reverse transcriptase from a group II intron of *Anoxybacillus flavithermus*

**DOI:** 10.1101/2023.12.09.570907

**Authors:** Igor P. Oscorbin, Maxim L. Filipenko

**Affiliations:** Institute of Chemical Biology and Fundamental Medicine, Siberian Branch of the Russian Academy of Sciences (ICBFM SB RAS), 8, Lavrentiev Avenue, Novosibirsk, 630090, Russia

**Keywords:** reverse transcriptase, group II intron, *Anoxybacillus flavithermus*, thermal stability, processivity, reverse transcription, RT-LAMP

## Abstract

Reverse transcriptases (RTs) are a family of enzymes synthesizing DNA using an RNA template and participating in retrovirus propagation and telomere lengthening. In vitro, RTs are widely applied in various methods, including RNA-seq, RT-PCR and RT-LAMP. Thermostable RTs from bacterial group II introns are promising tools for biotechnology; they showed higher thermostability, fidelity and processivity comparing to commonly using M-MuLV RT and its mutants. However, the diversity of group II intron-encoded RTs is still insufficiently studied. Here, we biochemically characterized a novel RT from a thermophilic bacterium Anoxybacillus flavithermus, isolated from a hot spring in New Zealand with an optimal temperature for growth around 60°C. The cloned RT, named Afl RT, retained around 40% of the specific activity after 45 min incubation at 50°C. Optimal pH was 8.5, temperature — 45-50 °C and Mn^2+^ ions were an optimal cofactor. In a processivity analysis with MS2 phage gRNA (3569 b), Afl RT elongated fully up to 36% of template molecules. In reverse transcription and RT-qLAMP, the enzyme allowed to detect up to 10 copies of MS2 phage genomic RNA per reaction. Thus, Afl RT is a promising enzyme for practical applications requiring thermostable processive RTs.

## 1 Introduction

Reverse transcriptases (RTs) comprise a specific family of DNA polymerases and synthetize DNA using RNA as a template. In eukaryotes, RTs take part in propagation of retroviruses, telomere maintained, and mobile genetic elements expansion. In prokaryotes, several classes of RTs mostly mediate various anti-phage immunity mechanisms. In vitro, RTs are indispensable tools for multiple practical applications that are used in research, bio-technology and molecular diagnostics.

Among multiple known RTs, the most commonly using enzyme is Moloney murine leukemia virus reverse transcriptase (M-MuLV RT). The enzyme is relatively well studied, and its mutated variants are available on a market with increased thermal stability, tolerance to inhibitors, processivity, etc. Other popular commercial RTs are avian leukemia virus RT (AVL RT) and HIV-1 RT that were also derived from retroviruses. However, not only retroviruses can be a source of reverse transcriptases; RTs were also found in prokaryotes. In last several decades, a number of bacterial RTs was identified, including enzymes from group II introns (GII introns) [1], retrons [2], diversity generating retroelements (DGR) [3], abortive bacteriophage infection (Abi) [4,5] and CRISPR-Cas systems [6]. With exception of GII introns, other genetic elements are involved in bacterial defense against phages, where mechanism of action assumes reverse transcription [6–9]. Exact functions of several other smaller RTs’ classes remain unknown. Around 40% of complete and draft genomes contain at least one gene, encoding putative RT. Thus, reverse transcriptases in prokaryotes are highly diverse, and their roles in cells still not completely uncovered.

In group II introns, RTs participate in propagation of mobile genetic elements by replicating their long coding sequences. GII-RTs demonstrated superior fidelity and processivity than retroviral RTs enabling more uniform coverage in RNA-seq than classical retroviral RTs [10,11]. Bacterial RTs also possess strong template-switching ability, beneficial for attachment of adaptors in RNA-seq and helps to omit RNA ligation or tailing steps. RTs from thermophilic bacteria are thermostable enzymes that allows to perform cDNA synthesis at temperatures around and above 50-60 °C. Higher reaction temperature destabilizes RNA secondary structures that can lead to RT stalling resulting in inefficient synthesis with short products. The latter is especially valuable for isothermal amplification; thermostable RTs can be used for single-tube analysis carried out in a constant single temperature. Thus, a simple and cost-effective point-of-care testing becomes possible. Taking in mind all mentioned benefits, RTs from bacterial group II introns, especially, from thermophilic bacteria are promising tools for practical application.

Currently, only scarce RTs or intron encoded proteins were reported from bacterial group II introns of Thermosynechococcus elongates [10], Geobacillus stearothermophilus [10,12], Eubacterium rectale [11], Oceanobacillus iheyensis [13], Lactococcus lactis [14], Sinorhizobium meliloti [15]. Among them, RTs from L. lactis, T. elongates, G. stearothermophilus and E. rectale were biochemically characterized and probed for practical applications. Thus, broad diversity of bacterial RTs remains practically unknown. In that sense, thermophilic bacteria can be a good source of new effective RTs.

Here, we cloned and characterized a novel reverse from a group II intron of a thermophilic bacteria Anoxybacillus flavithermus named Afl RT. The enzyme demonstrated optimal temperature around 45-50 °C and high processivity. Afl RT was also suitable for RT-qPCR and RT-qLAMP conferring sensitivity up to 10 MS2 phage genomic RNA molecules per reaction.

## 2. Materials and Methods

### 2.1 Search and selection of Afl RT coding sequence

For selection of reverse transcriptases genes, a tool MyRT was used from a work of Sharifi and Ye [16]. The tool allows to select putative genes of reverse transcriptases in all complete and bacterial genomes. The search was conducted in RTs of a GII class from draft and complete genomes, based on the presence of an RVT_1 domain (Pfam ID: PF00078) which is common for all known RTs. Found candidate GII-RTs were curated and only those from thermophilic hosts were further considered. At the last step, only RTs without any additional domains with exception of GIIM (maturases) were selected for possible cloning.

### 2.2 Expression and purification of Afl RT

The coding sequence of Afl RT was synthetized and cloned into pET28b vector by Shanghai RealGene Bio-tech, Inc. (Shanghai, China) resulting in a plasmid pAflRT.

A starter culture of E. coli BL21 (DE3) pLysS (Promega, WI, Madison, USA) strain harboring the plasmid pAflRT was grown to OD_600_ = 0.3 in LB medium with 25 μg/mL kanamycin at 37 °C. In a LiFlus GX fermenter (Biotron Inc., Bucheon, South Korea), 4 L of LB with 25 μg/mL kanamycin were inoculated with 40 ml of the starter culture, and the cells were grown to OD_600_ = 0.6 at 37 °C. Expression of Afl RT was induced by adding IPTG up to 1 mM concentration. After induction for 4 h at 37 °C, the cells were harvested by centrifugation at 4,000 × g and stored at −70 °C.

For a protein purification, the cell pellet was resuspended in a lysis buffer (50 mM Tris-HCl pH 9.0, 100 mM NaCl, 1 mM PMSF, 2 M urea, 1 mg/mL lysozyme), incubated for a 30 min on ice followed by sonication. After lysis, the soluble fraction was separated by two consequent centrifugation steps at 20,000×g for 30 min. Soluble proteins were precipitated ON by 60% (NH_4_)_2_SO_4_ followed by centrifugation at 20,000×g for 30 min. The resulting pellet was suspended in the lysis buffer and loaded onto a 5 ml IMAC column (Bio-Rad, CA, Hercules, USA) pre-equilibrated with a buffer A (50 mM Tris-HCl pH 8.0, 0.3 M NaCl), followed by washing the column with 25 ml of the buffer A with 1 M NaCl. Bound proteins were eluted using 10 column volumes 0-100% linear gradient of a buffer B (buffer A with 0.5 M imidazole). After the affinity chromatography, fractions with Afl RT were pooled and loaded onto a 2-ml Macro-Prep DEAE Resin (Bio-Rad, column CA, Hercules, USA) pre-equilibrated with a buffer C (50 mM Tris-HCl, 0.1 mM EDTA, pH 8.0). The column was washed with 10 ml of the buffer C, and bound proteins were eluted by a 10 column volumes 0–100% linear gradient of a buffer D (50 mM Tris-HCl, 1 M NaCl, 0.1 mM EDTA, pH 8.0). Fractions with Afl RT were pooled, dialyzed against a storage buffer (20 mM Tris-HCl, 200 mM KCl, 0.1 mM EDTA, 50% glycerol, pH 8.5) and stored at −20 °C. All fractions from each step were analysed by SDS-PAGE. Purity of the isolated Afl RT was not less than 95%. A concentration of purified Afl RT was measured using a standard Bradford assay.

### 2.3 Reverse transcriptase activity assay

The specific activity of Afl RT was assayed by using radiolabelled nucleotide incorporation. The reaction mix (50 µL) contained 0.4 mM poly(rA)/oligo(dT)25 (concentration defined by oligo(dT)42), 0.5 mM α-[32P]-dTTP (4 Bq/pmol), 50 mM Tris-HCI (pH 8.5), 6 mM MgCl_2_, 150 mM (NH_4_)_2_SO_4_, 5 mM DTT. The reactions were initiated by adding the enzyme on ice, with the samples immediately transferred to a preheated thermal cycler for incubation at 45 °C for 10 min, followed by inactivation by heating at 90 °C for a 1 min. The reaction products were collected on DE81 paper (Sigma-Aldrich, MO, St. Louis, USA), washed twice with 0.5 M Na_2_HPO_4_, and counted in a Pharos PX (Bio-Rad, CA, Hercules, USA). One unit of polymerase activity was defined as the amount of enzyme that incorporated 1 nmol of dTTP into acid-insoluble material in 10 min at 50 °C.

### 2.4 Thermal stability, optimal temperature and ion concentration

The thermal stability of Afl RT was studied by heating the enzyme followed by the assaying reverse transcriptase activity. The aliquots of the polymerase activity reaction buffer (described above) containing an identical amount 0.2 U of Afl RT were incubated at 40–80 °C by 10°C per step for 15-90 min. The reactions were chilled on ice, and the reverse transcriptase activity was measured as mentioned above.

The temperature optimum of Afl RT was defined by measuring reverse transcriptase activity at the range of 25–70°C by 5°C per step, while other conditions were identical to those described above for the reverse transcriptase activity assay.

The optimal ion concentrations were examined similarly as in the reverse transcriptase activity assay using 25 – 400 mM NH_4_Cl, KCl, 50 – 400 mM NaCl, Na_2_SO_4_, (NH_4_)_2_SO_4_ or 1 – 10 mM BaCl_2_, CaCl_2_, FeSO_4_, MgCl_2_, MnCl_2_, ZnSO_4_, CoCl_2_, CuCl_2_, NiCl_2_.

### 2.5 cDNA synthesis

The template RNA for cDNA synthesis was isolated from MS2 phage as described in the Supplementary. The 10 µL reaction probes containing MS2 RNA and 25 µM random N7 were heated for 3 min at 65°C, followed by cooling on ice at 5 min. The amount of MS2 RNA is specified for each experiment in the Results section. After priming, 10 µL of 2× reaction buffer (50 mM Tris-HCI (pH 8.5), 6 mM MgCl_2_, 150 mM (NH_4_)_2_SO_4_, 5 mM DTT) with 200 U of Afl RT were added to cooled template-primer mixtures, and the probes were immediately transferred to a preheated thermal cycler for 1 h incubation at 50 °C followed heating at 90°C for a 3 min. The reaction products were analysed using quantitative real-time PCR.

### 2.6 Quantitative PCR

Real-time RT PCR was performed in a CFX 96 thermocycler (Bio-Rad, Hercules, CA, USA) in a total reaction volume of 20 µL, containing a 1× PCR buffer (65 mM Tris-HCl pH 8.9, 24 mM (NH_4_)_2_SO_4_, 0.05% Tween 20, 2.5 mM MgCl_2_), 0.3 µM primers MS2-5-F/R and 0.1 µM probe MS2-5-P (Table 1), 1 unit of Taq-polymerase (Biosan, Novosibirsk, Russia) and a cDNA template as indicated in the Results section. The amplification was performed using the following program: denaturation at 95 °C for 3 min and 50 cycles with denaturation at 95 °C for 10 s, followed by annealing and elongation at 60 °C for 40 s with a registration of fluorescent signals in a FAM channel. Each experiment was conducted in three independent replicates, each run included a no-template control.

**Table 1.**
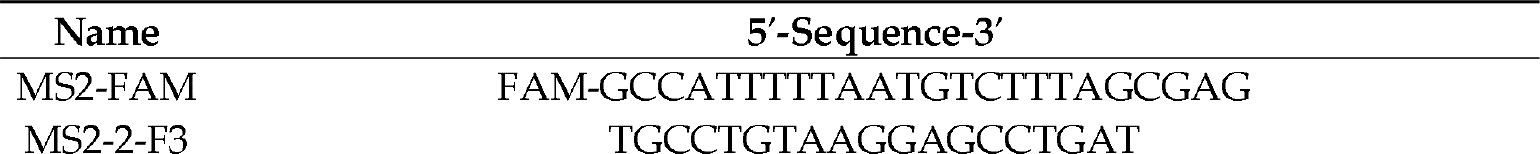
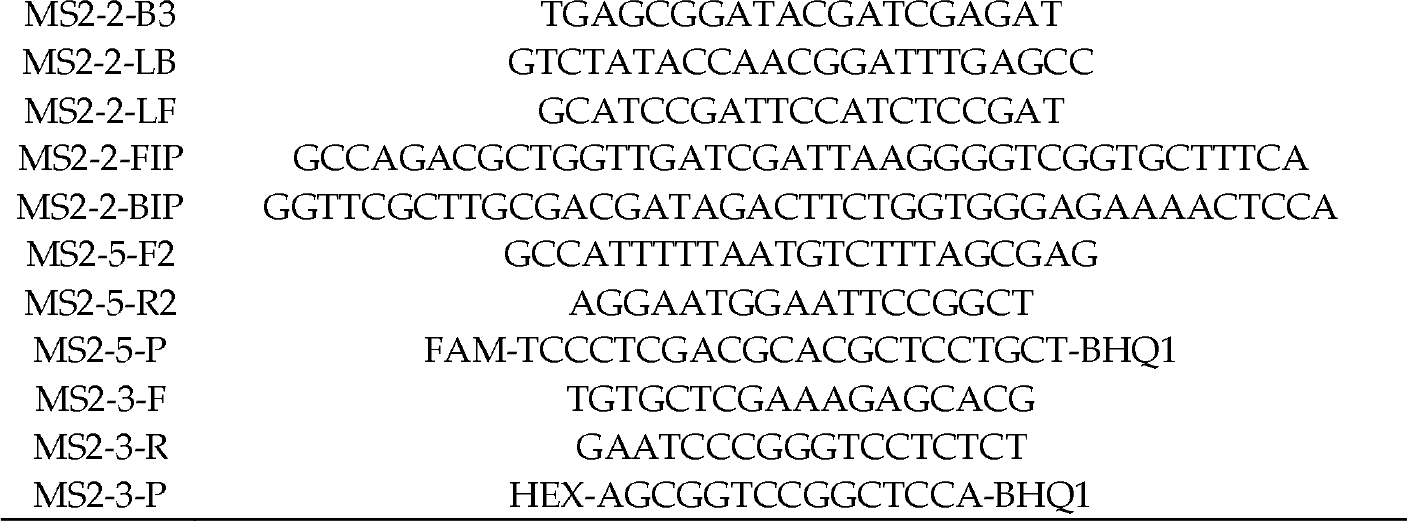
List of oligonucleotide primers and probes.

### 2.7. Reverse-Transcription Quantitative Loop-Mediated Isothermal Amplification (RT-qLAMP)

The reaction mixture for LAMP (20 µL) contained 1× reaction buffer for Bst-polymerase (20 mM Tris-HCl pH 8.8, 10 mM (NH_4_)_2_SO_4_, 50 mM KCl, 0.1% Tween-20, 8 mM MgSO_4_), 1.25 mM each dNTP, 0.4 µM each external primer (F3/B3), 0.8 µM loop primers (LF/BF), 1.6 µM internal primers (FIP/BIP) (Table 1), MS2 RNA as a template, 8 units of Gss-polymerase from Geobacillus sp. 777 [17], 200 U of Afl RT and 1 µM intercalating dye SYTO-82. The amount of MS2 RNA is specified for each experiment in the Results section. RT-LAMP was performed in the CFX96 thermocycler (Bio-Rad, CA, Hercules, USA) using a two-step program with the 15 min initial reverse transcription, followed by LAMP in a real-time mode. The exact temperature of the reverse transcription step is specified below in the Results section for each experiment. The program included the following steps: 90 cycles of primer annealing and elongation, each at 62 °C for 30 s with the registration of fluorescence signal in a HEX channel and post-amplification melting of amplification products in the range of 70–95 °C. Each experiment was conducted in three independent replicates, each run included a no-template control and a no reverse transcriptase control. Tt values (time-to-threshold, time interval before the intersection between an amplification curve and a threshold line) were calculated after each run and were used to assess RT-qLAMP efficacy.

### 2.8 Processivity assay

Reaction mixes in a 20 μL volume containing 5 ng MS2 RNA and 1 µM MS2-5-F2 primer (Table 1) were heated for 3 min at 65°C, followed by cooling on ice at 5 min. After priming, 10 µL of 2× reaction buffer (50 mM Tris-HCI (pH 8.5), 6 mM MgCl_2_, 150 mM (NH_4_)_2_SO_4_, 5 mM DTT) with 200 U of Afl RT were added to cooled template-primer mixtures, and the probes were immediately transferred to a preheated thermal cycler for 1 h incubation at 50 °C followed heating at 90°C for a 3 min. The reaction products were analysed using droplet digital PCR.

### 2.9 Droplet digital PCR

The ddPCR was performed using the QX100 system (Bio-Rad, Hercules, CA, USA) according to the manufacturer’s recommendations. A 20 µL ddPCR reaction mixture contained 1 × ddPCR master mix (Bio-Rad, Hercules, CA, USA), 0.9 µM primers, 0.25 µM probe (Table 1) and 2 mL of tested cDNA. The entire reaction mixture together with 70 µL of droplet generation oil (Bio-Rad, Hercules, CA, USA) was loaded into a disposable plastic cartridge (Bio-Rad, Hercules, CA, USA) and placed in a droplet generator. After processing, the droplets obtained from each sample were transferred to a 96-well PCR plate. The amplification was carried out using T100TM Thermal Cycler (Bio-Rad, Hercules, CA, USA) according to the program: DNA polymerase activation at 95 °C for 10 min followed by 45 cycles of PCR amplification (94 °C for 30 s and 58 °C for 60 s), and 98 °C for 10 min, 2 °C/s^−1^ ramp rate at all steps. After the PCR, the droplets were counted with the QX100 Droplet Reader. The data obtained were analyzed with QuantaSoft software (Bio-Rad, Hercules, CA, USA).

## 3. Results

### 3.1 Search of the Afl RT gene and purification of Afl RT

To search for a new RT from bacterial group II introns, we use an online tool published in a work of Sharifi and Ye [16]. The tool allows not only to predict novel RTs, but also search for putative reverse transcriptases among draft and complete genomes. We searched for novel GII RTs from thermophilic bacteria using the following criteria: presence of the common for RTs RVT_1 domain (Pfam ID: PF00078), thermophilic host and absence of any additional domains in an intron encoded protein with an exception of maturases GIIM. Thus, we tried to select putative thermostable RTs without any additional unpredictable enzymatic activities, which are often among bacterial reverse transcriptases. Following these criteria, among 6776 GII RTs from complete and 6209 GII RTs from draft genomes, we chose a single enzyme from group II intron of a thermophilic bacterium Anoxybacillus flavithermus. The host was found in hot spring in New Zealand with an optimal temperature for growth around 60°C [18]. Previously, several thermostable enzymes were isolated from A. flavithermus, including lipase [19], α-amylase [20], cyclomaltodextrinase [21], xylanases [22] with a temperature optimum in the range of 50-65 °C. Therefore, it was legitimate to assume the same thermostability of an RT from the same bacterium.

### 3.2 Biochemical properties of Afl RT

#### 3.2.1 Optimal temperature and thermostability

After purification, we studied biochemical properties of Afl RT, using fluorescently labelled oligo(dT)_40_ primer and poly(r)A template. Molar activity of Afl RT was 1.58*10^5^, which is close to molar activities of AVL RT and M-MuLV RT, reverse transcriptases from retroviruses [23].

High reaction temperature is beneficial for reverse transcription, melting complex RNA structures and increasing specificity of RTs against mismatched primers. To examine optimal reaction temperature for Afl RT, we carried out reverse transcription at 25-70 °C with a step of 5 °C and quantified reaction products. Results of optimal temperature assay are presented in Figure 1.

**Figure 1.**
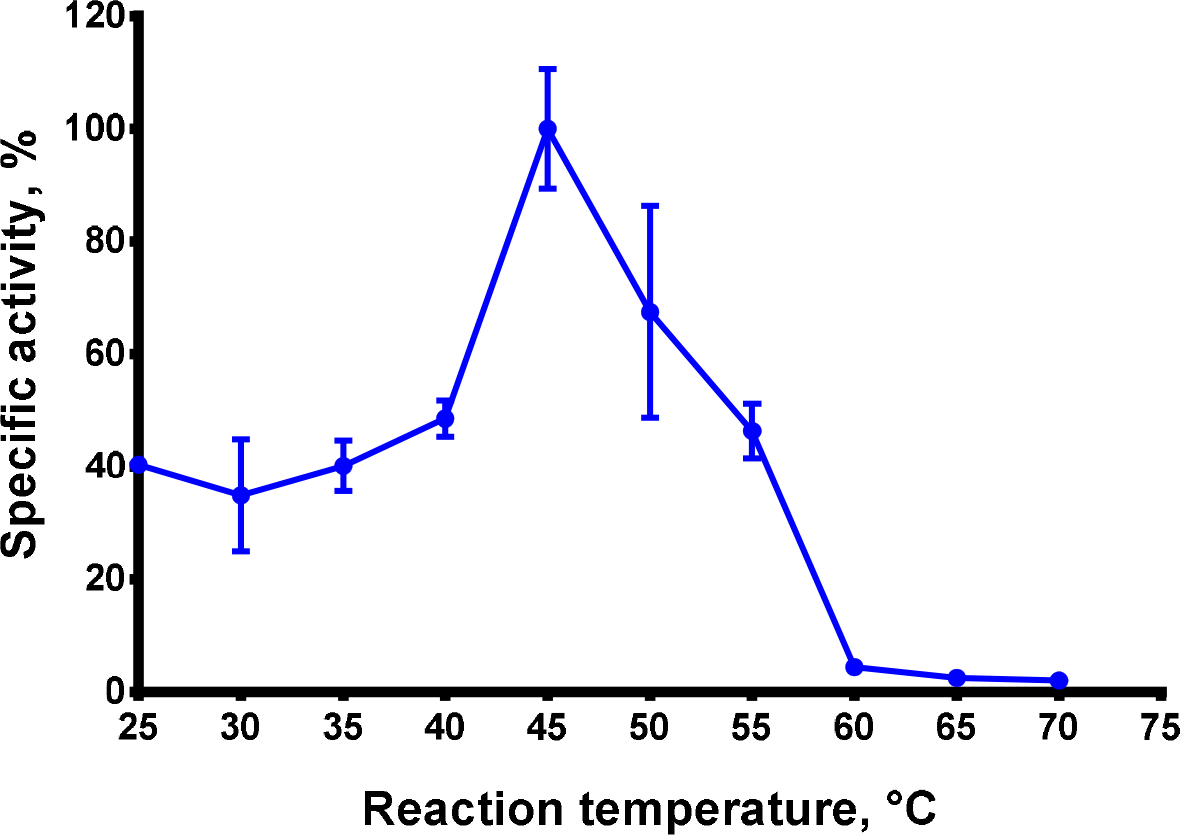
Optimal temperature assay. The optimal temperature for Afl RT was determined in RT activity assay using fluorescently labelled oligo(dT)_40_ primer in poly(rA)/oligo(dT)_40_ substrate. Each experiment was triplicated.

Afl RT demonstrated the highest reaction speed at 45-50 °C. This observation is a good agreement with optimal growth temperature of the host bacteria, Anoxybacillus flavithermus, with is around 60 °C. In further experiments, reverse transcription was conducted at 50 °C.

Next, we tested thermostability of Afl RT. Thermal stability is closely related with optimal reaction temperature and defines an ability of an enzyme to retain a specific activity after exposure to elevated temperatures. To examine thermostability of Afl RT, the enzyme was incubated at 40-80 °C for up to 90 min following RT activity analysis. Results of the assay are presented in Figure 2.

**Figure 2.**
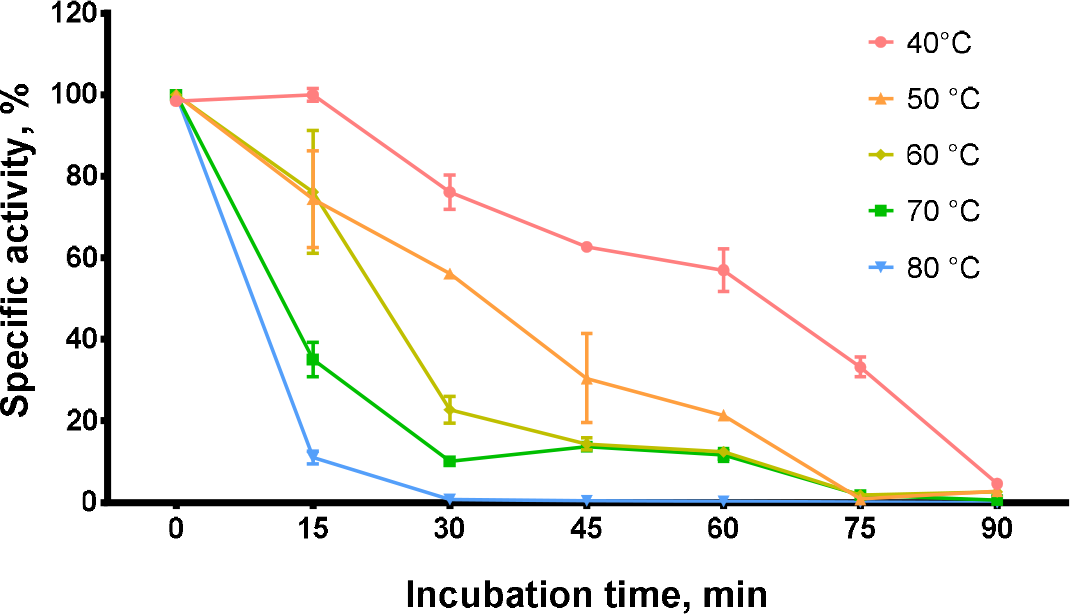
Thermostability assay. Afl RT was incubated at 40-80 °C for up to 90 min following RT activity assay using fluorescently labelled oligo(dT)_40_ primer in poly(rA)/oligo(dT)_40_ substrate. Each experiment was triplicated.

The enzyme retained 60% of RT activity after incubation at 50 °C for 30 minutes or 60 minutes at 40 °C and 40% — after 45 minutes at 50 °C. At higher temperatures, Afl RT lose the activity much faster: after 30 minutes at 70 °C only 10% of activity was detected and 30 minutes at 80 °C completely inactivated the reverse transcriptase. Thus, Afl RT was relatively thermostable comparing to M-MuLV RT.

#### 3.2.2 Optimal reaction buffer

A composition of reaction buffer, including pH, type and concentration of a buffer reagent, salts, detergents and other additives such as DTT, defines the efficacy of enzymatic catalysis. Ionic strength and pH effect on a conformation and interaction of the enzyme and its substrate, which leads to favorable or unfavorable condition for chemical transformations in an active site of the enzyme. Another important factor are cofactors, e.g. divalent cations in the case of reverse transcriptases. To devise an optimal reaction buffer for Afl RT, we titrated several salts at various pH in reverse transcription following selection of a type and concentration of an optimal divalent cation. A concentration of buffer reagent was similar in all experiments: 50 mM Tris-HCl for pH 7.0-9.5 and 25 mM MES for pH 6.0-6.5. For divalent cations, a concentration of dNTP was fixed as 0.5 mM. Results of titration experiments are presented in Figure 3.

**Figure 3.**
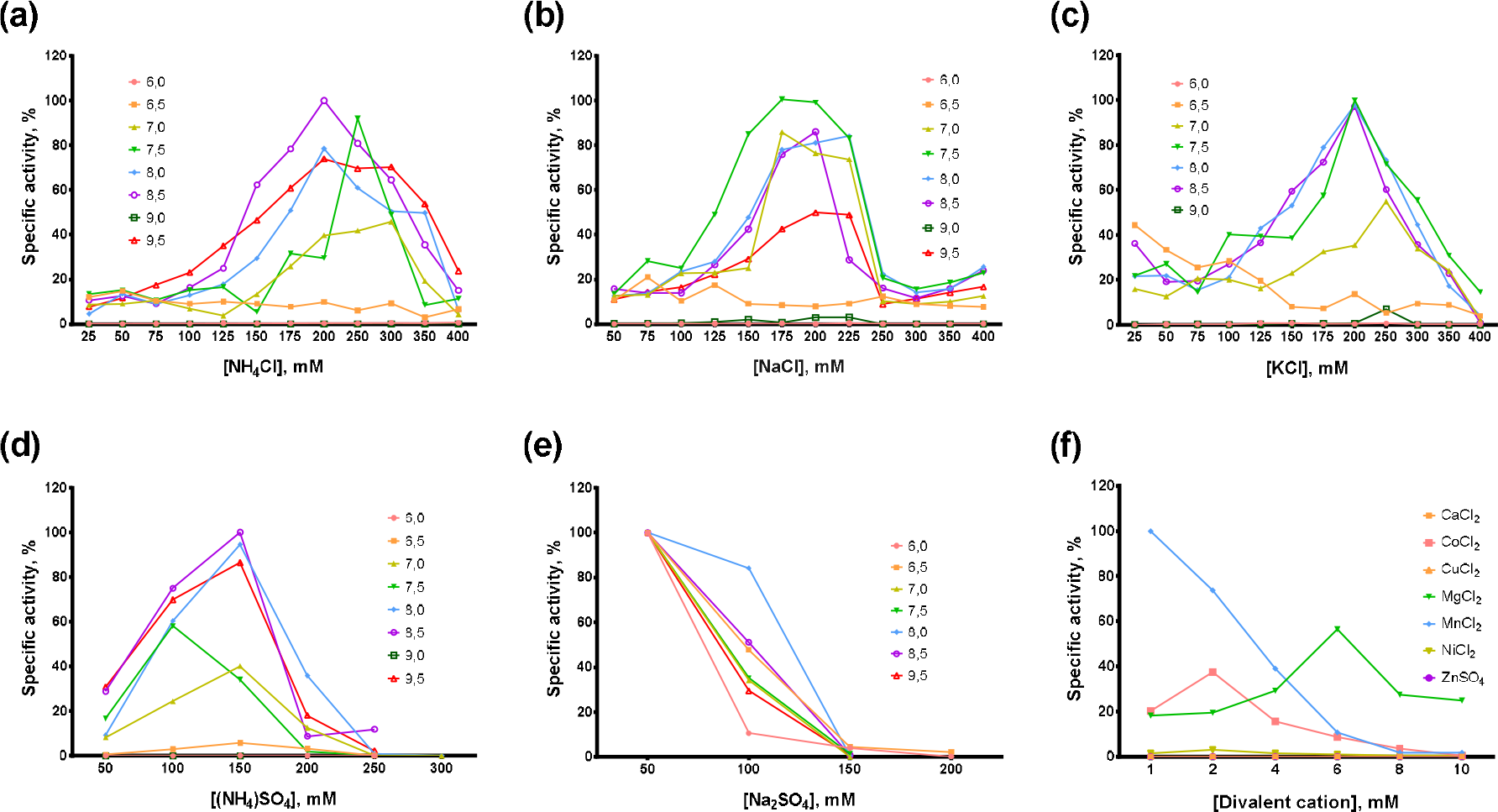
Optimal reaction buffer for Afl RT. Salts concentration and pH were varied in RT activity assay with fluorescently labelled oligo(dT)_40_ primer in poly(rA)/oligo(dT)_40_ substrate: a) NH_4_Cl, b) NaCl, c) (NH_4_)_2_SO_4_, d) Na_2_SO_4_, e) divalent cations Ba^2+^, Ca^2+^, Co^2+^, Cu^2+^, Fe^2+^, Mg^2+^, Mn^2+^, Ni^2+^, Zn^2+^.

In most cases, optimal salt concentration was independent from pH of the reaction buffer. Only for NH_4_Cl, KCl and pH 7.0, the optimal salt concentration was 50-100 mM higher than for other pH values. Optimal concentration of sulfates was lower than of chlorides: 200-250 mM NH_4_Cl and 150 mM (NH_4_)_2_SO_4_, 200-250 mM NaCl and 50 mM Na_2_SO_4_, respectively. K_2_SO_4_ was not titrated because of high ionic strength and relatively low solubility of this sulfate. When different salts were compared, the highest activity of Afl RT was observed with 150 mM (NH_4_)_2_SO_4_ and pH 8.5 which was used in all following experiments.

Among divalent cations, Afl RT was completely inactive with CuCl_2_, ZnSO_4_. With CaCl_2_, only residual activity was observed, less than 1% of the specific activity with MgCl_2_; an inhibition effect of NiCl_2_ was less striking and Afl RT retained around 5% of activity comparing to MgCl_2_. Despite low activity with NiCl_2_ and total inhibition by FeSO_4_, Afl RT was surprisingly active with CoCl_2_ — activity with 2 mM CoCl_2_ achieved 66% activity with 6 mM MgCl_2_. The most favorable cofactor for Afl RT was MnCl_2_ instead of MgCl_2_, and the enzyme was 1.75 folds more active with MnCl_2_. However, in following experiments we used 6 mM MgCl_2_ due to higher error rate of synthesis with Mn^2+^ cations which was shown for other RTs [24–26]. Also, Afl RT synthetized more cDNA in the presence of 5 mM DTT (data not shown).

### 3.3 Afl RT in practical applications

From a practical angle, it is important to assess RTs in various applications. Novel enzymes could be more efficient than popular commercial reverse transcriptases and become valuable tools for modern biotechnology. Here, we chose reverse transcription, RT-PCR and RT-LAMP as the most common approaches with RTs. MS2 page genomic RNA served as a template and was titrated in the range 1*10^2^-1*10^8^ copies per reaction.

#### 2.3.1 Afl RT in reverse transcription

In reverse transcription, 200 U of Afl RT was used per reaction with random N7 primers, reaction mixes were incubated at 50 °C for 1 hour. After reverse transcription, amount of cDNA was quantified by qPCR and plotted in Figure 4.

**Figure 4.**
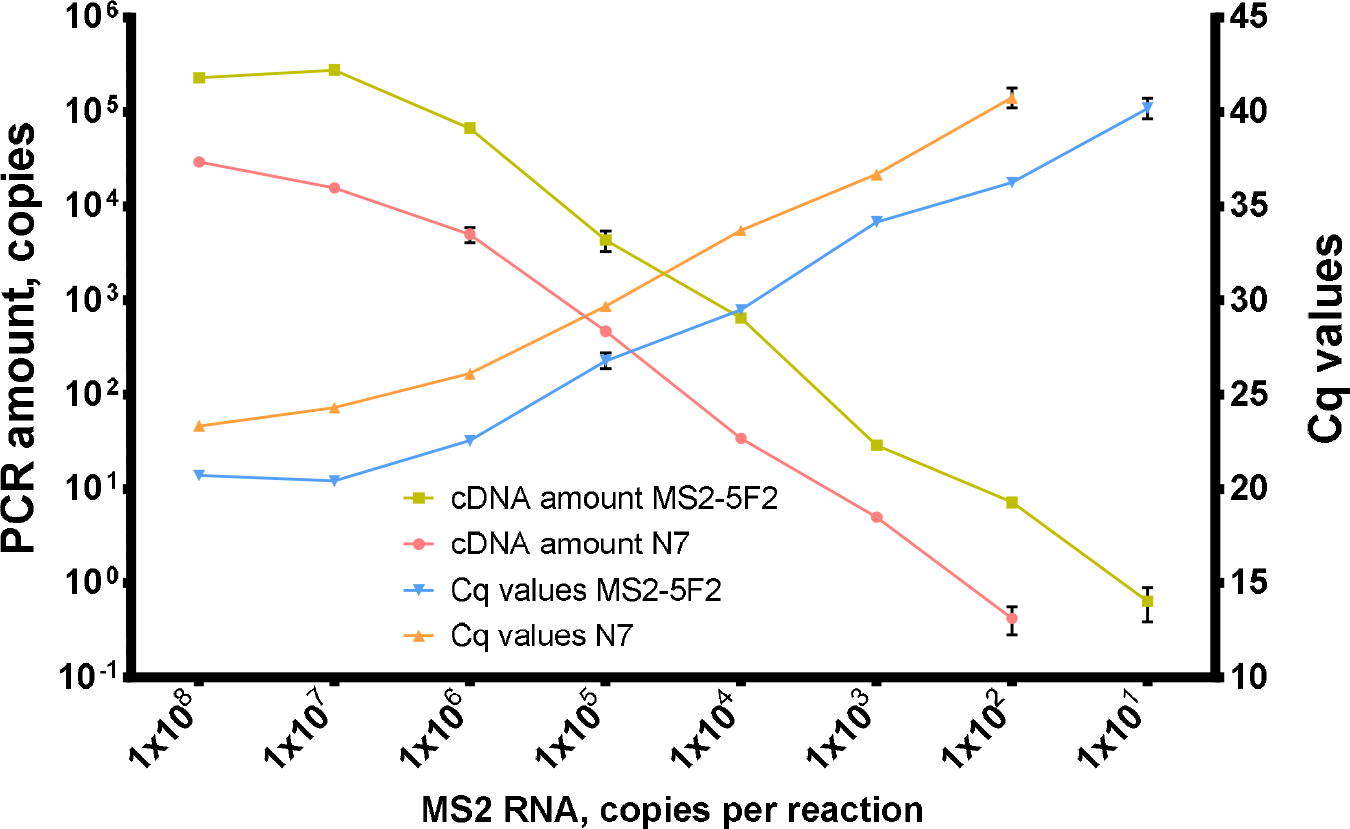
Titration of MS2 genomic RNA in reverse transcription with Afl RT. Reverse transcription was conducted with 1*10^2^-1*10^8^ of MS2 genomic RNA as template at 50 °C for 40 minutes with random hexamers and 200 U of Afl RT per reaction. Reaction products were quantified by qPCR. cDNA amount and Cq values were plotted against template concentration. Each experiment was triplicated.

With N7 primers, Afl RT successfully synthetized cDNA when down to 1*10^2^ copies of MS2 genomic RNA was added in reverse transcription mixes. With a specific MS2-5F2 primer, reverse transcription was around 10 times more effective: more cDNA was produced with the same amount of the template and the least detectable template concentration was 10^1^ copies of MS2 genomic RNA per reaction. Plausibly, the observed higher efficacy of reverse transcription with the specific primer could be a result of its higher melting temperature. N7 primers are much shorter and most of them could not anneal at 50 °C.

#### 2.3.2 Afl RT in RT-qLAMP

In RT-qLAMP, 200 U of Afl RT was used per reaction. The reverse transcription step was conducted at 40, 50 or 60 °C. Sequences for LAMP primers MS2-F3/B3/LB/LF/FIP/BIP are given in Table 1. With each temperature and template concentration, a control reaction was performed without Afl RT to test reverse transcriptase activity of Gss-polymerase. When RT-qLAMP ended, LAMP products were examined by melting curves analyzes. Resulted time-to-threshold (Tt) values are presented in Figure 5.

**Figure 5.**
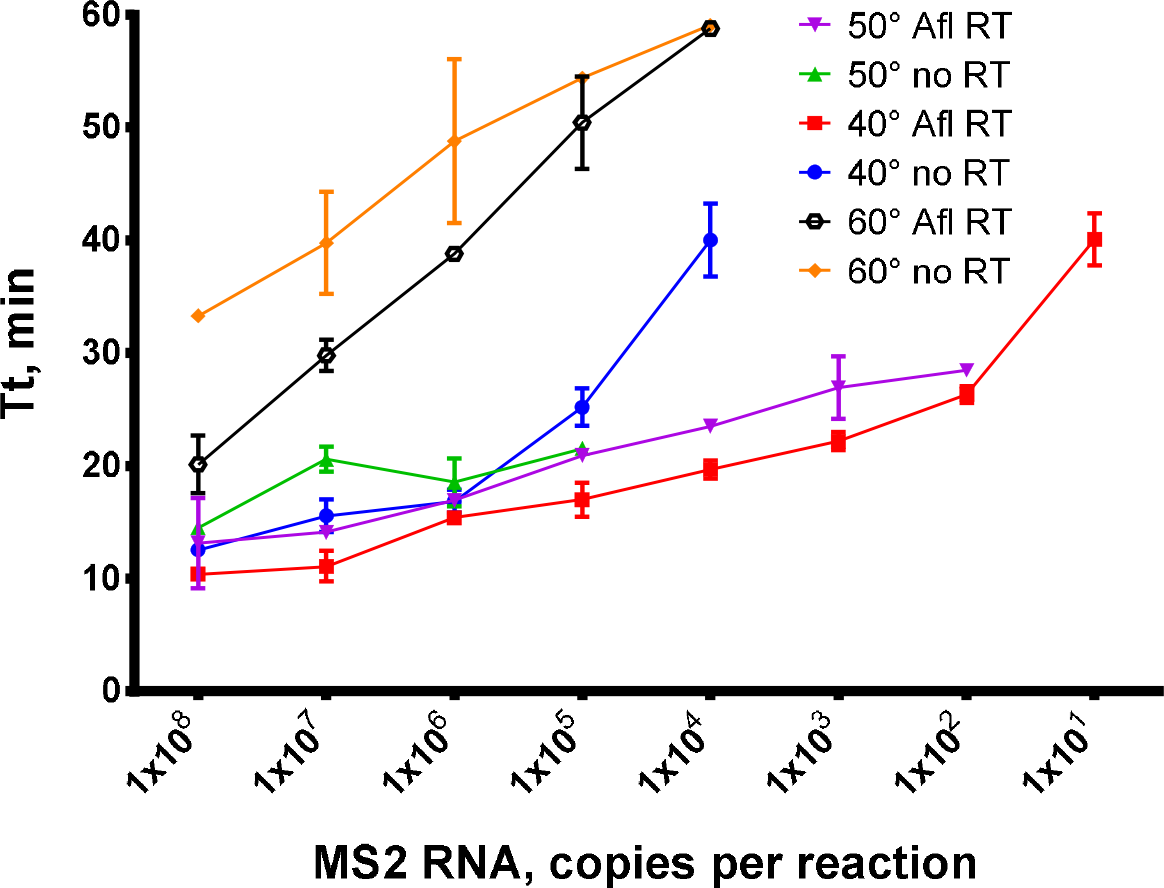
Titration of MS2 genomic RNA in RT-qLAMP with Afl RT. RT-qLAMP was conducted with 0.5 fg-500 ng pf MS2 genomic RNA as template and 200 U of Afl RT per reaction. The temperature of a reverse transcription step was 40, 50, 60 °C for 15 minutes. Time-to-threshold values were plotted against a template concentration. Each experiment was triplicated.

Similar with reverse transcription, addition of Afl RT in RT-qLAMP allowed to detect up to 10^1^ copies of MS2 genomic RNA when a temperature of reverse transcription was 40 °C, and 10^2^ copies per reaction when the temperature was 50 °C. Without a specific reverse transcriptase, RT-qLAMP was much less sensitive, 10^5^ at 50 °C and 10^4^ at 40 °C, indicating a weak reverse transcriptase ability of Bst-like Gss-polymerase. Efficacy of reverse transcription at 60 °C was much lower, which can be a result of Afl RT thermal inactivation. However, despite loss of activity, 10^4^ copies of MS2 genomic RNA were successfully detected. Thus, the most effective reverse transcription in RT-qLAMP was at 40-50°C.

#### 2.3.3 Processivity of Afl RT

Processivity is an important parameter of reverse transcriptases, defining their ability to synthetize full-length cDNA fragments. Short partially elongated cDNA could lead to issues in further analysis, especially NGS. In that sense, a special case are GC-rich highly structured RNA molecules, and MS2 genomic RNA is a good model of such complex templates. Here, we studied processivity of Afl RT using MS2 genomic RNA (3569 nt) as a template for long cDNA synthesis. The length of cDNA products was assessed using droplet digital PCR with two sets of primers and TaqMan probes for both 5′- and 3′-ends of the cDNA (Table 1). The 5′/3′-ends ratio served as a surrogate marker of Afl RT processivity: a processive enzyme that synthetizing longer cDNA molecules should have a 5′/3′-ends ratio closer to 100%. The results of processivity assay are presented in Figure 6.

**Figure 6.**
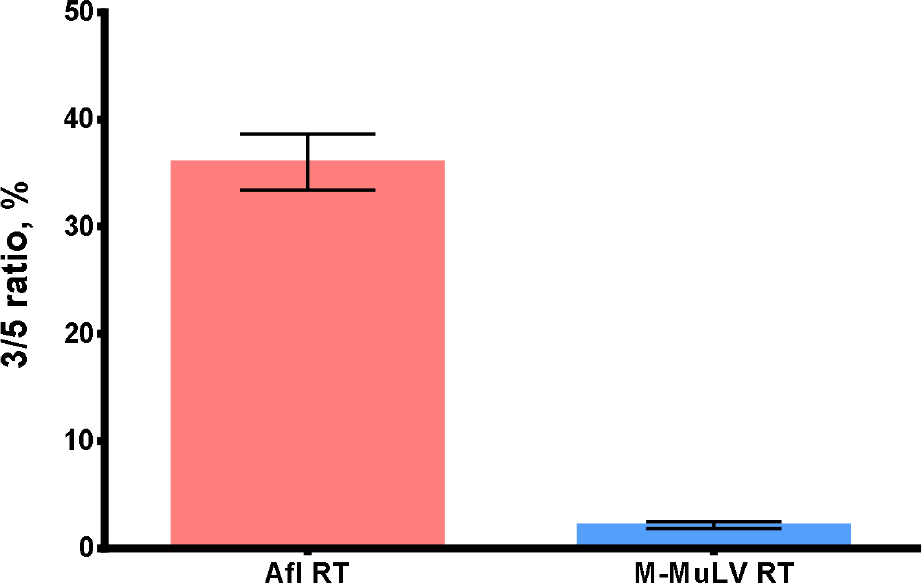
The 5′/3′-ratio of cDNA synthetized by Afl RT and M-MuLV RT. MS2 genomic RNA (3569 nt) served as a template for reverse transcription. Each reaction was run in three technical repeats. The 5′/3′-ratio of cDNA ends was determined using ddPCR with primers for 5′- and 3′-ends of MS2 genomic RNA.

The 5′/3′-ratio of cDNA ends for Afl RT was close to 36.0 %, indicating 1/3 of all cDNA molecules being full-length, which was significantly higher than 2.2 % for M-MuLV RT. Thus, processivity of Afl RT several folds exceeded M-MuLV RT and was similar to other GII reverse transcriptases.

## 4. Discussion

Reverse transcriptases were firstly discovered in retroviruses and retroviral RTs became model enzymes for studies of RNA conversion to cDNA. Needless to say, retroviral RTs have also become essential tools in regards of RNA studies, requiring conversion of RNA to more stable and convenient cDNA. Among all retroviral RTs, the most common commercial enzyme is monomeric M-MuLV RT and its mutants with improved thermal stability, processivity, and tolerance to inhibitors. Dimeric AMV RT and HIV-1 RT are more thermostable than wild-type M-MuLV RT, yet, all these enzymes are inactivated by incubation for several minutes at temperatures higher than 70-75 °C [27–29]. Processivity and fidelity of retroviral RTs are lacking, being in the range 14-97 b and 1 mismatch per 10^4^ incorporated nucleotides, respectively [30,31]. These parameters limit usage of common RTs for practical applications where uniform precise cDNA synthesis is crucial, e.g. RNA-seq, especially, in a single-cell format. Short cDNA molecules lead to underrepresentation of reads corresponding to 5’-ends of mRNA molecules. Mismatches and other altercations in reads, such as indels, increase the noise and decrease sensitivity for rare events. For these reasons, novel more thermostable and accurate RTs are in a high demand.

While RTs from retroviruses are relatively well-studied and introduced in biotechnology, reverse transcriptases from prokaryotes still remain a dark matter. In the latest phylogenic analysis, prokaryotic RTs were divided to 42 classes, and functions of only 11 classes have been unearthed [16]. For instance, discovered 30 years ago retrons were shown to participate in anti-phage defense only in 2020 [7]. This lack of knowledge becomes striking comparing to DNA polymerases from other families participating in replication and DNA repair. Thus, reverse transcriptases wait for systematical biochemical studies to reveal in vivo role of these enzymes.

Among bacterial RTs, the most abundant are enzymes from type II introns or GII RTs encompassing around 65 % of all putative RTs [16]. Simultaneously, several GII RTs were cloned, purified and biochemically characterized which provided two exceptionally processive, precise and relatively thermostable RTs from T. elongates (TGIRT) and E. rectale (Marathon RT) [10,11]. However, this is insufficient for systematical comparison of RTs from type II introns with retroviral counterparts. Also, some properties of GII RTs remained undescribed including optimal reaction buffer that is crucial for the best performance in practical applications.

Here, we focused on GII RTs from thermophilic bacteria, as our main goal was to find novel enzymes with a potential for biotechnology applications. For that reason, we excluded RTs with any additional domains and took into consideration only enzymes with a classical domain structure with fingers, palm, and an X-domain (thumb) [32]. A GII RT from a thermophilic bacterium Anoxybacillus flavithermus met all mentioned criteria and was chosen for the further biochemical analysis.

GII RTs are notoriously known for their insolubility, problems with purification and a low yield after isolation [10–12,33]. In bacteria, the enzymes exist as ribonucleoproteins, where a protein part tightly binds the respective coding RNA. Also, GII RTs bear a high positive charge and have an isoelectric point close to 9-10. This is believed to result in a tendency to form insoluble conglomerates during a heterologous expression in E. coli cells. In early attempts to purify GII RTs, the enzymes lose greatly the specific activity and were highly contaminated by E. coli proteins [12,33]. Expression at 18-20 °C often allowing to obtain proteins is a soluble fraction, also was unsuccessful. Later, the insolubility problem was partially solved by fusion of GII RTs either with a maltose-binding MalE protein via a rigid alanine linker in TGIRT [10] or with a SUMO tag in Marathon RT [11]. Interestingly, RTs fused with MalE via a common flexible linker were still insoluble. Here, we managed to purify Alf RT without linking it with any solubility tags using only a 2 M urea as a mild chaotropic agent. Plausibly, slight disruption of interactions with RNA molecules, proteins and other cell components was enough for purification of a GII RT without fusion with helper solubulity domains. However, this discrepancy of our results with observations of other groups needs to be clarified specifically, as GII RTs have similar pI, net charge and domain structure.

In previous works describing GII RTs, authors mostly aimed to demonstrate superior thermostability, processivity and template switching activity of these enzymes. A proper description of these enzymatic properties is necessary for practical application. However, an important issue for practical applications is optimization of a reaction buffer. Suboptimal conditions could decrease a reaction yield and the length of cDNA products. Here, we firstly studied relative activity of a GII RT in the presence of various salts and divalent cations. Interestingly, Afl RT was more active in the presence of Mn^2+^ than with conventional Mg^2+^. To our knowledge, it was not reported before for any other GII RT and similar effect was noted before for retroviral M-MuLV RT [34]. Fidelity of DNA polymerases and RTs is usually lower with uncanonical cofactors [24,35]. Thus, Mg^2+^ ions are preferable for correct synthesis, especially, taking in account a possible application of GII RTs in single-cell RNA-seq.

As other characterized GII RTs, Afl RT demonstrated high processivity being able to fully elongate 1/3 of all cDNA molecules using as a template MS2 genomic RNA. Seemingly, processive synthesis is a characteristic feature of all GII RTs that is necessary for propagation of group II introns in cells. The same goes for thermostability, enzymes from moderate thermophiles living at 50-60 °C are relatively thermostable in vitro too. While optimal reaction temperature for Afl RT was lower than for TGIRT and Marathon RT, RT-LAMP with Afl RT demonstrated a good sensitivity in the optimal buffer for Gss-polymerase. Thus, Afl RT can become an alternative for common commercial retroviral RTs.

Several limitations of the present study need to be mentioned. Direct comparison of a new Afl RT with TGIRT and Marathon RT would provide more information about the link between the structure and enzymatic properties of GII RTs. GII RTs are also known for their strong terminal transferase activity, which was not tested for Afl RT. Suitability of Afl RT for RT-PCR also needs to be assessed, and we plan to perform corresponding experiments in the future.

To sum up, we cloned a new reverse transcriptase from a group II intron of a thermophilic bacterium. Anoxybacillus flavithermus. The novel enzyme, Afl RT was biochemically characterized and tested in reverse transcription and RT-LAMP uncovering its potential for biotechnology.

## 5. Conclusions

In this work, we biochemically characterized and tested in practical applications a novel RT from a thermophilic bacterium Anoxybacillus flavithermus. The cloned Afl RT was relatively thermostable comparing to retroviral RTs and showed higher processivity. The enzyme also allowed to detect up to 10 copies of an RNA template in reverse transcription and RT-LAMP. Thus, Afl RT is a promising enzyme for biotechnology.

## Author Contributions

Conceptualization, M.L.F.; methodology, I.P.O.; validation, I.P.O. and M.L.F.; formal analysis, I.P.O.; investigation, I.P.O.; data curation, I.P.O.; writing—original draft preparation, I.P.O.; writing—review and editing, M.L.F.; visualization, I.P.O.; supervision, M.L.F.; project administration, I.P.O.; funding acquisition, M.L.F. All authors have read and agreed to the published version of the manuscript.

## Institutional Review Board Statement

Not applicable.

## Informed Consent Statement

Not applicable.

## Data Availability Statement

The data presented in this study are available on request from the corresponding author.

## Disclaimer/Publisher’s Note

The statements, opinions and data contained in all publications are solely those of the individual author(s) and contributor(s) and not of MDPI and/or the editor(s). MDPI and/or the editor(s) disclaim responsibility for any injury to people or property resulting from any ideas, methods, instructions or products referred to in the content.

## References

1. Michel, F.; Ferat, J.-L. Structure and Activities of Group II Introns. Annu. Rev. Biochem. 1995, 64, 435–461, doi:10.1146/annurev.bi.64.070195.002251.

2. Lampson, B.C.; Inouye, M.; Inouye, S. Reverse Transcriptase with Concomitant Ribonuclease H Activity in the Cell-Free Synthesis of Branched RNA-Linked MsDNA of Myxococcus Xanthus. Cell 1989, 56, 701–707, doi:10.1016/0092-8674(89)90592-8.

3. Liu, M.; Deora, R.; Doulatov, S.R.; Gingery, M.; Eiserling, F.A.; Preston, A.; Maskell, D.J.; Simons, R.W.; Cotter, P.A.; Parkhill, J.; et al. Reverse Transcriptase-Mediated Tropism Switching in Bordetella Bacteriophage. Science (80-.). 2002, 295, 2091–2094, doi:10.1126/science.1067467.

4. Fortier, L.-C.; Bouchard, J.D.; Moineau, S. Expression and Site-Directed Mutagenesis of the Lactococcal Abortive Phage Infection Protein AbiK. J. Bacteriol. 2005, 187, 3721–3730, doi:10.1128/JB.187.11.3721-3730.2005.

5. Wang, C.; Villion, M.; Semper, C.; Coros, C.; Moineau, S.; Zimmerly, S. A Reverse Transcriptase-Related Protein Mediates Phage Resistance and Polymerizes Untemplated DNA in Vitro. Nucleic Acids Res. 2011, 39, 7620–7629, doi:10.1093/nar/gkr397.

6. Silas, S.; Mohr, G.; Sidote, D.J.; Markham, L.M.; Sanchez-Amat, A.; Bhaya, D.; Lambowitz, A.M.; Fire, A.Z. Direct CRISPR Spacer Acquisition from RNA by a Natural Reverse Transcriptase-Cas1 Fusion Protein. Science 2016, 351, aad4234, doi:10.1126/science.aad4234.

7. Millman, A.; Bernheim, A.; Stokar-Avihail, A.; Fedorenko, T.; Voichek, M.; Leavitt, A.; Oppenheimer-Shaanan, Y.; Sorek, R. Bacterial Retrons Function In Anti-Phage Defense. Cell 2020, 183, 1551–1561.e12, doi:10.1016/j.cell.2020.09.065.

8. Mestre, M.R.; Gao, L.A.; Shah, S.A.; López-Beltrán, A.; González-Delgado, A.; Martínez-Abarca, F.; Iranzo, J.; Redrejo-Rodríguez, M.; Zhang, F.; Toro, N. UG/Abi: A Highly Diverse Family of Prokaryotic Reverse Transcriptases Associated with Defense Functions. Nucleic Acids Res. 2022, 50, 6084–6101, doi:10.1093/nar/gkac467.

9. Guo, H.; Arambula, D.; Ghosh, P.; Miller, J.F. Diversity-Generating Retroelements in Phage and Bacterial Genomes. Microbiol. Spectr. 2014, 2, doi:10.1128/microbiolspec.MDNA3-0029-2014.

10. Mohr, S.; Ghanem, E.; Smith, W.; Sheeter, D.; Qin, Y.; King, O.; Polioudakis, D.; Iyer, V.R.; Hunicke-Smith, S.; Swamy, S.; et al. Thermostable Group II Intron Reverse Transcriptase Fusion Proteins and Their Use in CDNA Synthesis and Next-Generation RNA Sequencing. RNA 2013, 19, 958–970, doi:10.1261/rna.039743.113.

11. Zhao, C.; Liu, F.; Pyle, A.M. An Ultraprocessive, Accurate Reverse Transcriptase Encoded by a Metazoan Group II Intron. RNA 2018, 24, 183–195, doi:10.1261/rna.063479.117.

12. Vellore, J.; Moretz, S.E.; Lampson, B.C. A Group II Intron-Type Open Reading Frame from the Thermophile Bacillus (Geobacillus) Stearothermophilus Encodes a Heat-Stable Reverse Transcriptase. Appl. Environ. Microbiol. 2004, 70, 7140–7147, doi:10.1128/AEM.70.12.7140-7147.2004.

13. Toor, N.; Keating, K.S.; Taylor, S.D.; Pyle, A.M. Crystal Structure of a Self-Spliced Group II Intron. Science 2008, 320, 77–82, doi:10.1126/science.1153803.

14. Matsuura, M.; Saldanha, R.; Ma, H.; Wank, H.; Yang, J.; Mohr, G.; Cavanagh, S.; Dunny, G.M.; Belfort, M.; Lambowitz, A.M. A Bacterial Group II Intron Encoding Reverse Transcriptase, Maturase, and DNA Endonuclease Activities: Biochemical Demonstration of Maturase Activity and Insertion of New Genetic Information within the Intron. Genes Dev. 1997, 11, 2910–2924, doi:10.1101/gad.11.21.2910.

15. Muñoz-Adelantado, E.; San Filippo, J.; Martínez-Abarca, F.; García-Rodríguez, F.M.; Lambowitz, A.M.; Toro, N. Mobility of the Sinorhizobium Meliloti Group II Intron RmInt1 Occurs by Reverse Splicing into DNA, but Requires an Unknown Reverse Transcriptase Priming Mechanism. J. Mol. Biol. 2003, 327, 931–943, doi:10.1016/s0022-2836(03)00208-0.

16. Sharifi, F.; Ye, Y. Identification and Classification of Reverse Transcriptases in Bacterial Genomes and Metagenomes. Nucleic Acids Res. 2022, 50, e29–e29, doi:10.1093/nar/gkab1207.

17. Oscorbin, I.P.; Boyarskikh, U.A.; Filipenko, M.L. Large Fragment of DNA Polymerase I from Geobacillus Sp. 777: Cloning and Comparison with DNA Polymerases I in Practical Applications. Mol. Biotechnol. 2015, 57, 947–959, doi:10.1007/s12033-015-9886-x.

18. Heinen, W.; Lauwers, A.M.; Mulders, J.W. Bacillus Flavothermus, a Newly Isolated Facultative Thermophile. Antonie Van Leeuwenhoek 1982, 48, 265–272, doi:10.1007/BF00400386.

19. Bakir, Z.B.; Metin, K. Purification and Characterization of an Alkali-Thermostable Lipase from Thermophilic Anoxybacillus Flavithermus HBB 134. J. Microbiol. Biotechnol. 2016, 26, 1087–1097, doi:10.4014/jmb.1512.12056.

20. Özdemir, S.; Matpan, F.; Okumus, V.; Dündar, A.; Ulutas, M.S.; Kumru, M. Isolation of a Thermophilic Anoxybacillus Flavithermus Sp. Nov. and Production of Thermostable α-Amylase under Solid-State Fermentation (SSF). Ann. Microbiol. 2012, 62, 1367–1375, doi:10.1007/s13213-011-0385-4.

21. Aliakbari, N.; Mirzaee, Z.; Jafarian, V.; Khalifeh, K.; Salehi, M. Genetic and Biochemical Characterization of a Novel Thermostable Cyclomaltodextrinase From Anoxybacillus Flavithermus. Starch - Stärke 2019, 71, doi:10.1002/star.201800133.

22. Ellis, J.T.; Magnuson, T.S. Thermostable and Alkalistable Xylanases Produced by the Thermophilic Bacterium Anoxybacillus Flavithermus TWXYL3. ISRN Microbiol. 2012, 2012, 1–8, doi:10.5402/2012/517524.

23. Yasukawa, K.; Nemoto, D.; Inouye, K. Comparison of the Thermal Stabilities of Reverse Transcriptases from Avian Myeloblastosis Virus and Moloney Murine Leukaemia Virus. J. Biochem. 2008, 143, 261–268, doi:10.1093/jb/mvm217.

24. Kristen, M.; Plehn, J.; Marchand, V.; Friedland, K.; Motorin, Y.; Helm, M.; Werner, S. Manganese Ions Individually Alter the Reverse Transcription Signature of Modified Ribonucleosides. Genes (Basel). 2020, 11, doi:10.3390/genes11080950.

25. Valverde-Garduño, V.; Gariglio, P.; Gutiérrez, L. Functional Analysis of HIV-1 Reverse Transcriptase Motif C: Site-Directed Mutagenesis and Metal Cation Interaction. J. Mol. Evol. 1998, 47, 73–80, doi:10.1007/PL00006364.

26. Sirover, M.A.; Loeb, L.A. On the Fidelity of DNA Replication. Effect of Metal Activators during Synthesis with Avian Myeloblastosis Virus DNA Polymerase. J. Biol. Chem. 1977, 252, 3605–3610.

27. Malboeuf, C.M.; Isaacs, S.J.; Tran, N.H.; Kim, B. Thermal Effects on Reverse Transcription: Improvement of Accuracy and Processivity in CDNA Synthesis. Biotechniques 2001, 30, 1074–1084, doi:10.2144/01305rr06.

28. Konishi, A.; Yasukawa, K.; Inouye, K. Improving the Thermal Stability of Avian Myeloblastosis Virus Reverse Transcriptase α-Subunit by Site-Directed Mutagenesis. Biotechnol. Lett. 2012, 34, 1209–1215, doi:10.1007/s10529-012-0904-9.

29. Álvarez, M.; Matamoros, T.; Menéndez-Arias, L. Increased Thermostability and Fidelity of DNA Synthesis of Wild-Type and Mutant HIV-1 Group O Reverse Transcriptases. J. Mol. Biol. 2009, 392, 872–884, doi:10.1016/j.jmb.2009.07.081.

30. DeStefano, J.J.; Buiser, R.G.; Mallaber, L.M.; Myers, T.W.; Bambara, R.A.; Fay, P.J. Polymerization and RNase H Activities of the Reverse Transcriptases from Avian Myeloblastosis, Human Immunodeficiency, and Moloney Murine Leukemia Viruses Are Functionally Uncoupled. J. Biol. Chem. 1991, 266, 7423–7431, doi:10.1016/S0021-9258(20)89464-2.

31. Potapov, V.; Fu, X.; Dai, N.; Corrêa, I.R.; Tanner, N.A.; Ong, J.L. Base Modifications Affecting RNA Polymerase and Reverse Transcriptase Fidelity. Nucleic Acids Res. 2018, 46, 5753–5763, doi:10.1093/nar/gky341.

32. Lambowitz, A.M.; Zimmerly, S. Group II Introns: Mobile Ribozymes That Invade DNA. Cold Spring Harb. Perspect. Biol. 2011, 3, a003616–a003616, doi:10.1101/cshperspect.a003616.

33. Ng, B.; Nayak, S.; Gibbs, M.D.; Lee, J.; Bergquist, P.L. Reverse Transcriptases: Intron-Encoded Proteins Found in Thermophilic Bacteria. Gene 2007, 393, 137–144, doi:10.1016/j.gene.2007.02.003.

34. Roth, M.J.; Tanese, N.; Goff, S.P. Purification and Characterization of Murine Retroviral Reverse Transcriptase Expressed in Escherichia Coli. J. Biol. Chem. 1985, 260, 9326–9335.

35. Vashishtha, A.K.; Konigsberg, W.H. The Effect of Different Divalent Cations on the Kinetics and Fidelity of Bacillus Stearothermophilus DNA Polymerase. AIMS Biophys. 2018, 5, 125–143, doi:10.3934/biophy.2018.2.125.

