## Supplementary for "A novel thermostable and processive reverse transcriptase from a group II intron of *Anoxybacillus flavithermus*": Supplementary materials.docx

**1.2. Isolation of MS2 Phage RNA**

The MS2 phage (GenBank ID NC_001417) was grown using the modified protocol of Sambrook and Russel [1]. The fresh night culture of *E. coli* K12 strain was diluted in 3 mL of MS2 medium to the OD_600_ = 1 (1×10^9^ cells/mL), followed by the addition of MS2 phage to reach the phage/cell ratio of 5. The cultures were incubated at 37 °С for 20 min, mixed with 500 mL of preheated MS2 medium, and incubated under the same conditions for 12 h. The chloroform was added to lyse the cells, and the culture was vortexed for 10 min at 37 °С. The lysate was treated by DNase I and RNase A (50 mg/mL each) for 30 min at 37 °С. Then, NaCl was added to the final concentration of 1 M. The mixture was incubated on ice for 1 h, with the debris separated by centrifugation (10 min, 11000×g) at 4 °С. The supernatant was supplied by the additional amount of ammonium sulfate to the final concentration of 50% (m/m), the mixture was incubated for 2 h at 4 °С, and the phage particles were precipitated by centrifugation (30 min, 11000×g) at 4 °С. The phage particles precipitated were dissolved in 30 mL of the TSM buffer (20 mM Tris-HCl, pH 7.4, 150 mM NaCl, 2 mM CaCl_2_, 2 mM MgCl_2_). MS2 RNA was isolated from phage particles using QIAamp Circulating Nucleic Acid Kit (Qiagen, Venlo, Netherlands) according to the manufacturer’s protocol and stored at −80 °С.

**1.3. Droplet Digital PCR**

The concentration of MS2 RNA was measured by a digital PCR using the QX200™ Droplet Digital™ PCR System (Bio-Rad, CA, Hercules, USA) according to the manufacturer’s instructions. The reactions were carried out in a total volume of 20 µL containing the DNA under examination (approximately 10^3^ copies per 20 µL), One-Step RT-ddPCR kit (Bio-Rad, CA, Hercules, USA), 900 nM primers MS2-5-F/R and 250 nM probe MS2-5-P (Table 1). For droplet generation, 20 µL of the PCR mix and 70 µL of the droplet generation oil were placed into corresponding wells of the DG8 cartridge, and the droplets were obtained in a droplet generator. Then, 40 µL of the obtained droplets were transferred to the 96-well PCR plate, foil-sealed, and placed into the thermocycler. The amplification was performed using the following program: reverse transcription at 42 °C for 60 min, enzyme activation at 95 °С for 10 min, followed by 45 cycles of 95 °С for 30 s, 58 °С for 60 s, with final heating for 10 min at 98 °С. The ramp rate was 2 °С/s for all steps. The droplets were analyzed by the droplet reader, and the data obtained were processed by the QuantaSoft package (Bio-Rad, CA, Hercules, USA).

***References***

1. Evans GA. Molecular cloning: A laboratory manual. Second edition. Volumes 1, 2, and 3. Current protocols in molecular biology. Volumes 1 and 2 [Internet]. Cell. New York: Cold Spring Harbor Laboratory Press; 1990. https://doi.org/10.1016/0092-8674(90)90210-6
